# py_ped_sim - A flexible forward genetic simulator for complex family pedigree analysis

**DOI:** 10.1101/2024.03.25.586501

**Authors:** Miguel Guardado, Cynthia Perez, Shalom Jackson, Joaquín Magaña, Sthen Campana, Emily Samperio, Berenice Chavez Rojas, Selena Hernandez, Kaela Syas, Ryan Hernandez, Elena I. Zavala, Rori Rohlfs

## Abstract

**Background:** Large-scale family pedigrees are commonly used across medical, evolutionary, and forensic genetics. These pedigrees are tools for identifying genetic disorders, tracking evolutionary patterns, and establishing familial relationships via forensic genetic identification. However, there is a lack of software to accurately simulate different pedigree structures along with genomes corresponding to those individuals in a family pedigree. This limits simulation-based evaluations of methods that use pedigrees.

**Results:** We have developed a python command-line-based tool called py_ped_sim that facilitates the simulation of pedigree structures and the genomes of individuals in a pedigree. py_ped_sim represents pedigrees as directed acyclic graphs, enabling conversion between standard pedigree formats and integration with the forward population genetic simulator, SLiM. Notably, py_ped_sim allows the simulation of varying numbers of offspring for a set of parents, with the capacity to shift the distribution of sibship sizes over generations. We additionally add simulations for events of misattributed paternity, which offers a way to simulate half-sibling relationships. We validated the accuracy of our software by simulating genomes onto diverse family pedigree structures, showing that the estimated kinship coefficients closely approximated expected values.

**Conclusions:** py_ped_sim is a user-friendly and open-source solution for simulating pedigree structures and conducting pedigree genome simulations. It empowers medical, forensic, and evolutionary genetics researchers to gain deeper insights into the dynamics of genetic inheritance and relatedness within families.

## Background

Genetic pedigrees are essential for studying medical, evolutionary, and forensic genetic inheritance. Pedigrees have been instrumental in understanding disease prevalence within families and populations and guiding genetic counseling and management strategies. They have been used to unravel the genetic influence of psychiatric disorders and neurodegenerative diseases via family-based studies **[1,2,3]**. Additionally, pedigrees help give us insight into rare variants acting on disease **[2,4,5,6]**. Recent efforts have shown that sharing rare variants is related to individuals’ demographic and familial history **[7]**, emphasizing the need to understand how rare variations are transmitted through families. In evolutionary genetics, family pedigree studies provide insights into population dynamics **[8,9,10]**, the heritability of traits **[11, 12, 13]**, and natural selection **[14, 15]**. The emerging forensic practice of investigative genetic genealogy (IGG) relies heavily on pedigree analysis to connect genetic relatives **[16, 17, 18]**. By constructing pedigrees and tracing familial relationships, investigators can identify potential suspects and narrow down a pool of individuals who may be related to unidentified remains and DNA evidence found at a crime scene **[19, 20]**.

Many pedigrees lack complementing genetic data; to overcome this, genetic simulations are leveraged to generate individual genomes based on available pedigree data. Current software exists for simulating genomes onto family pedigrees in various ways, such as sim1000G **[21]**, RarePedSim **[22]**, SimRVSequences **[23]**, and Msprime **[24]**. Current software is restricted to simulating a constrained number of variants due to computational complexity or is limited to simulating exons for the medical use of acting causal variants from family-based studies. Furthermore, these tools overlook crucial elements of evolutionary theory within their simulation framework, such as considering recombination and mutation rates. They are often overlooked due to the added complexity for users to implement. Current evolutionary genetic simulation approaches use either a forward or coalescent model. Coalescent simulations, such as those Msprime utilizes to simulate genomic pedigrees through a fixed ancestry model **[25]**, work backward, starting with present-day individuals and tracing their ancestral lineages to the most recent common ancestor (MRCA). This approach is practical when simulating genomes across larger time scales.

On the other hand, forward simulations are individual-based and progress downwards from the top of the pedigree, simulating one generation at a time until reaching the end. SLiM is a popular forward evolutionary simulator that can simulate genomes based on fixed pedigrees **[26, 27]**. This approach can offer more customizable initialization of founders within the pedigree and initialize founders easily with empirical genomes. While SLiM has a feature to simulate genomes based on fixed family pedigrees, it requires prior knowledge of the family structure, such as the list of founders in the pedigree and the generation in which descendants are created. Having to identify this information manually causes a barrier to performing genomic simulations on large sets of family pedigrees in an automated manner.

Automating the identification of this information alone by inputting a pedigree alone would create more accessibility to performing forward evolutionary simulations of the pedigree.

While the tools mentioned above can be used to simulate varying types of DNA profiles onto a set pedigree, there are still challenges in generating diverse and realistic pedigree structures. One way to help bridge this gap is through simulating pedigree structures. It is important to consider complex and realistic pedigree structures to accurately represent genetic diversity and the evolutionary history of a family lineage, giving us more insight into an individual’s genetic inheritance [**28**]. A limitation to simulating pedigree structures is the ability to simulate misattributed genetic paternity (MAP). A different parent mistakenly attributes the child’s biological paternity during this process. While it has been widely accepted that these MAP events can occur in 10% of our population [**29**], a meta-analysis suggested this number may be overestimated for most populations [**30**]. Modeling these events into the pedigree structure can help us understand inconsistency within family pedigrees and the impact that can have on the analysis of genetic kinship. Additionally, we must pay attention to how sibship size varies over generation and geography. It has long been acknowledged that sibship size has decreased in the recent century. Furthermore, the rate of that decrease changed dramatically across different countries [**31**], needing to explore how resulting family pedigree structures change across different geographical rates of sibship. Having the ability to use real-world data to simulate pedigree structures will allow open avenues to explore questions of how genetic variation develops across diverse demographic histories. However, no software exists to simulate realistic pedigrees that incorporate nonuniform rates of sibship across generations or generate non-paternity events introducing half-relationships.

Having a tool that enables the simulation of genomes onto realistic pedigree structures would open up new avenues of genetics research. By incorporating realistic pedigree structures into genomic simulations, we can better reflect the complexities of human populations and improve our understanding of how demographic history impacts the genetic variation within kin. Here, we present py_ped_sim, a command line tool in Python designed to facilitate genetic pedigree analysis. Our software incorporates four key features (Figure 1a): (1) the simulation of varied genetic pedigree structures based on sibship sizes over time, (2) the simulation of misattributed paternity events within family pedigrees, (3) the simulation of genomes based on fixed family pedigree, and (4) the identification of pairwise relationships within a given family pedigree (**Figure 1a**). To enable efficient genetic simulations on complex or incomplete pedigrees, we have developed a wrapper for SLiM that allows users to input varied pedigree data. We additionally demonstrate that py_ped_sim accurately generates genomes on family structures with expected genetic relationships.

**Figure 1:**
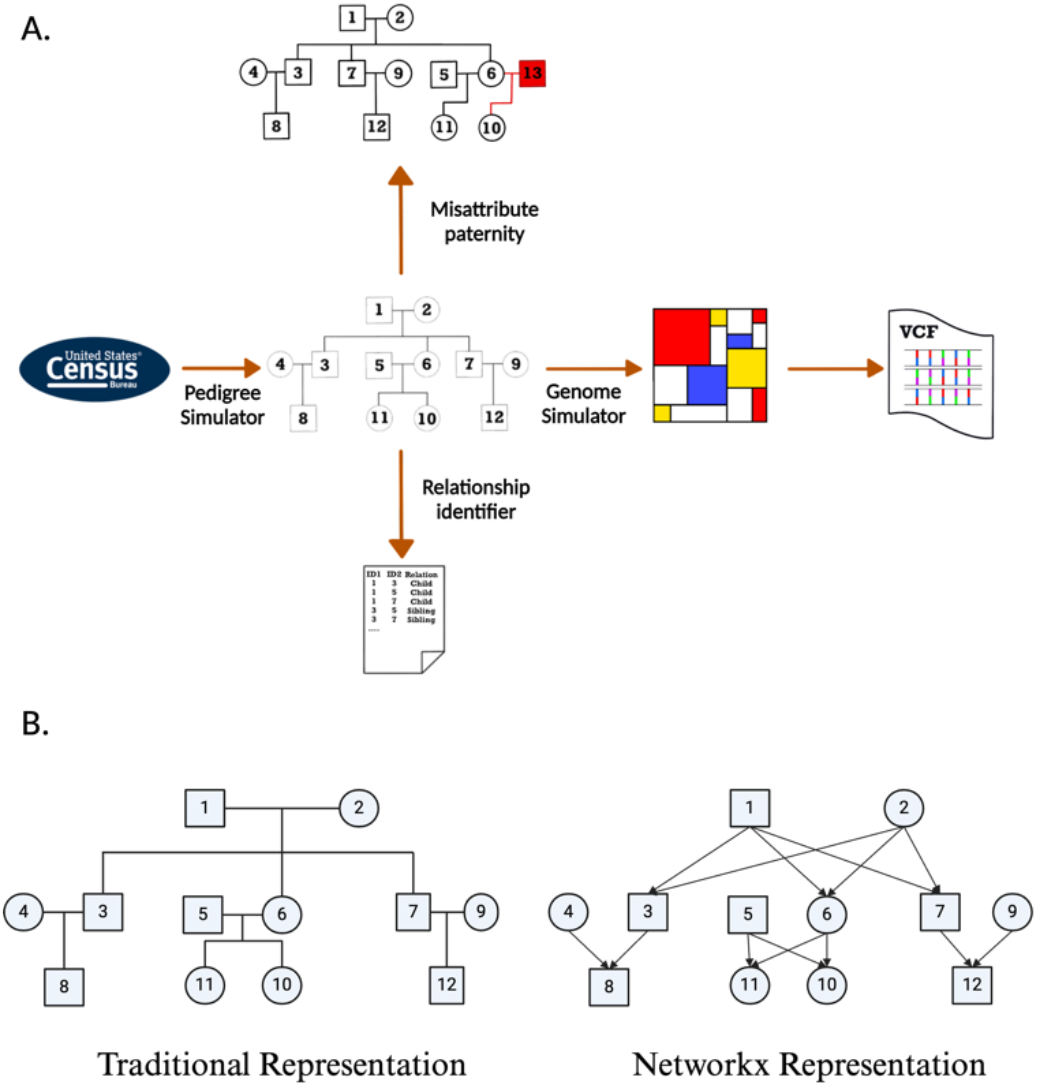
A) Visual abstract illustrating the primary features of the py_ped_sim software. Comparison of data structures used to represent family pedigrees. Our software utilises networks represented pedigrees.

### Implementation

**Definitions [Fig 2, Pedigree Schematic] (To Be Included as a Table):**

- DAG - Directed Acyclic Graph. Where individuals are nodes and edges are parent-child relationships.
- Descendant - offspring; an individual that has at least one known parent.
- Generation Tick - We define the generation of an individual the same way as we do in the SLiM simulation framework. Generations are a biological unit of time, having to do with things like the lifespan of an organism, the mean age at first reproduction, and so forth.
- Root Founder - A node that is connected to all lower-level nodes. The root founder of the pedigree will have a generation time of one.
- Explicit Founders - Individuals specified in the pedigree with no known ancestors.
- Implicit founders - Individuals in the pedigree with only one known ancestor. The implicit founder is considered the missing parent inside the family pedigree.
- Pedigree formats
- Traditional pedigree formal (.ped/.fam): This data representation of a family is traditionally used where the columns represented as [FID, IID, P1, P2, Sex, Phenotype]
- Networkx-based pedigree (.nx): Family pedigree based on a directed acyclic graph implemented in networkx. The files are two-column files where the first column is the parent and the second is a descendant.
- Slim-Readable pedigree (_slim_pedigree.txt): Family pedigree file that is readable by SLiM. Similar to a .ped file but includes and orders the generation in which descendants are created [Gen, IID, P1, P2].

**Figure 2:**
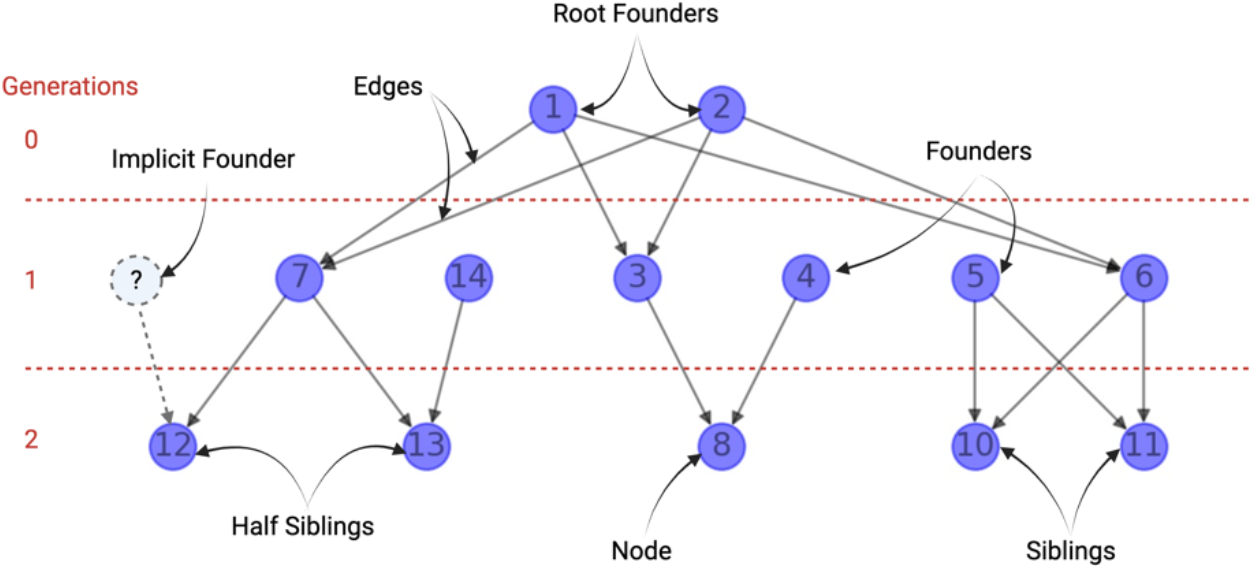
Schematic Overview of the pedigree definition used in our software.

### Dependencies and Versions

This software is developed in python (version 3.88), using SLiM (version 3.6), and bcftools (version 1.8). All user interaction with the software is through a Python front end. All software dependencies can be accessed via a conda virtual environment.

### Data Structure of Family Pedigree

We represent family pedigrees as directed acyclic graphs (DAGs) using the networkx package in python 3.8 **[32]**. The DAG comprises of nodes and edges, where nodes represent individuals and directed edges indicate parent-to-child genetic transmission **[Figure 1b, networkx pedigree schematic]**. Because an individual cannot be their genetic ancestor in a sexual reproductive system, these graphs are acyclic, meaning the directed path of the edges never forms a loop.

### Pedigree Simulator

py_ped_sim forward simulates pedigrees with the number of offspring per pair of genetic parents taken from user-provided data on sibship sizes. The novelty of our approach is the ability to vary the sibship distributions used across generations. We draw sibship sizes from normal distribution based on the user-provided generation, mean, and standard deviation. The user will include a csv file on each year’s mean and standard deviation of sibship size. For convenience, we provide a default sibship distribution file with data from the United States Census obtained via IPUMs **[33]**.

We utilize a depth-first recursive approach to simulate children until the last generation is reached. Users specify the number of generations in the pedigree via the number of generation ID requested. The number of offspring is drawn from user-provided sibship sizes for each set of parents for the generation number. Our software additionally simulates an individual’s sex and keeps track of the generation time in which individuals are created. We define the sex of an individual by whether a fertile individual produces a sperm (XY) or an egg (XX). It is important to note that we don’t simulate sex chromosomes in our simulation framework. We simulate the sex of an individual via drawing from a Bernoulli distribution to determine the sex of the first parent and set the sex of the second parent to the complement.

The output of py_ped_sim is a pedigree in networkx format, in addition to a profile file that contains the sex and generation time of everyone simulated. py_ped_sim pedigree output as DAGs via networkx format files (.nx).

### Misattributed Paternity Simulator

Modeling MAP events can offer another way to simulate more dynamic family pedigrees and investigate how MAP can impact downstream conclusions. We have incorporated a feature for simulating misattributed paternity events on existing family pedigrees. A complimentary benefit of introducing this to family pedigree simulations is adding genetic half relationships.

Our simulations of MAP events involve a two-step process: identifying individuals with MAP, descendants where MAP events will happen, and identifying the correct genetic parent where the new individual will be generated. First, we must identify individuals with whom MAP will happen, we will iterate across all individuals with specified genetic parents in the pedigree and simulate a binary event via a Bernoulli distribution. When identifying an individual where a MAP event will occur, we determine if the new parent will come from within an existing father in family pedigree, or if we will create a new individual to be the father. If we choose to use an existing parent, we sample randomly from fathers of the same generation tick as the father being replaced. If no potential fathers can be used within the same generation tick, py_ped_sim will automatically generate a new father to perform the MAP.

py_ped_sim, takes as input a networkx family pedigree file and a profiles file specifying sex and generational tick for individuals. Additionally, the user can input two parameters that specify the probability rate of a MAP event and whether the newly assigned genetic parent will be within the family, or to create a new individual. The output is the updated networkx family pedigree file and profile files, including newly created individuals.

### Genome Simulator

We introduce a feature to use SLiM to simulate genomic variation onto family pedigrees. This feature is a wrapper script that extracts features of the family pedigrees that SLiM requires for genomic simulations. Both explicit and implicit founders are to be identified alongside generation numbers for all descendants created. py_ped_sim represents the pedigree as a DAG; in that format, we can extract information on descendants’ generation time and founders of the pedigree. The output is a vcf file of the simulated genomes for everyone in the pedigree. The vcf does not include the genomes of implicit founders, that is, parents not specified in the original pedigree.

To initiate the genomic simulations, users specify the genetic pedigree structure, along with a genomic file (.vcf) for initializing the founders of the family. The user can specify the mutation and recombination rates for the simulations, default values have a mutation rate of 1e-8 and recomb at 1e-7. We provide two methods for initializing the founder genomes based on a user-provided vcf file. The first method randomly assigns each founder to an individual in the vcf file. Alternatively, users can provide an additional text file that assigns specific founders to corresponding IDs in the vcf file, offering more control over the initialization process. The founder genome can be simulated as a supplementary feature, which is particularly useful if an appropriate genome dataset for founder initialization is unavailable. The simulation is a neutral burn in simulations where you can specify the size of the simulated populations and how many individuals you want to sample from the population.

Identifying founders in simulations is crucial to supplying genomes for individuals without identified parents in the pedigree, establishing the genetic variation for individuals for all non-descendants. We identify individuals with no predecessors (explicit founders) and one known predecessor (implicit founder events). We start by identifying the family’s root founders and assigning them a generation time of one. The generation times of descendants are then assigned based on their shortest path to root founders. For individuals not directly descended from a root founder, we calculate the generational gap between the individual in question and a root founder’s descendant and use the difference to determine the generation number for the individual in question. In the event of consanguinity, when individuals have more than one path to the root founder, we ensure the child’s generation number is after their parent’s.

### Pairwise Relationships Identifier

py_ped_sim provides a systemic and quantified representation of relationships for all pairs of individuals inside a pedigree structure. Our software can quantify genetic relationships between pairs of individuals in a pedigree, defined by three metrics: meiotic distance (MD), the generation depth difference (GDD), and the genetic relationship type (GRT) **[Figure 3]**. These three statistics help us to code pairwise genetic relationships into genetic relationship categories (siblings, half-first cousins, etc.).

**Figure 3.**
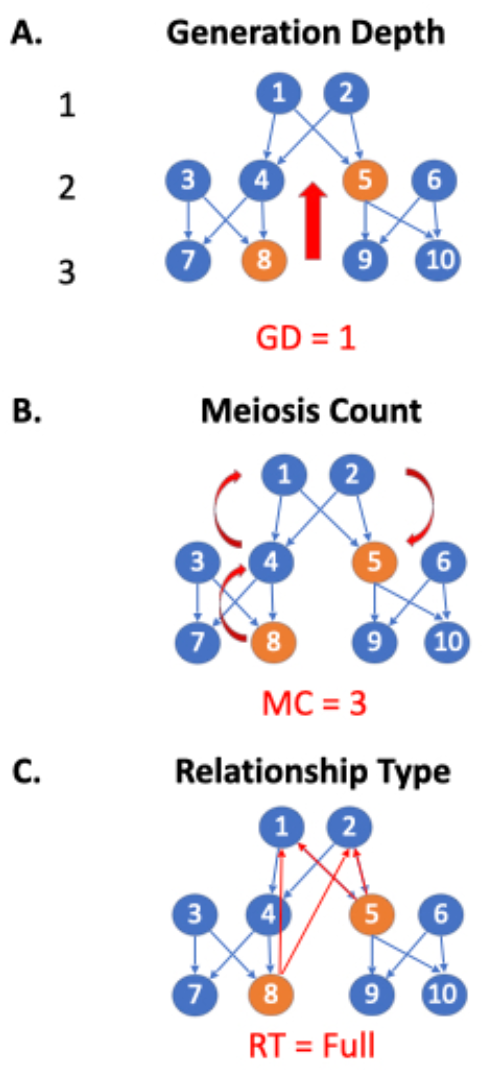
Visual representation of how each relationship metric in an avuncular relationship is determined. (A) Generation depth determines how many generations ticks separate two individuals; for an avuncular relationship, only one generation separates a child from their uncle. (B) Meiosis count is the shortest path for two relationships, building a path from their common ancestors. For an avuncular relationship, their common ancestor is the child’s grandparent and avuncular parents, producing a meiosis count of 3. (C) Relationship type determines if the relationship is a direct common ancestor, full, or half relationship. For this avuncular relationship shown, they share two common ancestors, determining this a full relationship.

GRT describes if two individuals are full genetic relatives (two shared ancestors), half genetic relatives (one shared ancestor), direct genetic relatives (defined above), or not determinable (NA). Let’s take the example of an avuncular relationship. If there are two common ancestors for a child and their uncle/aunt, the child’s grandparents and the parent of the avuncular relationship, then the relationship is classified as a full relationship **[Fig 3a]**. To determine GRT, we find all the shortest paths between two nodes. To determine direct and half relatives, we count the number of shortest paths two individuals are connected. If there is one path, they’re half-genetic relatives; if there are two paths, they’re full genetic relates.

Finally, the GDD between two individuals is the number of generations separating them. Using the same examples of avuncular and half-sibling relationships, we see GDDs of 1 and 0, respectively **[Fig 3b]**. GDD is the shortest path between the two nodes for individuals with a direct genetic connection.

Otherwise, GDD equals the absolute value of the difference between the shortest path lengths from the two individuals to their shared ancestor.

The meiotic distance between two individuals describes how much meiosis separates them. An avuncular (ex: aunt-nephew) relationship has an MC of three **[Fig 3c]**. Half siblings have an MC of two since they connect via one shared parent. If a direct genetic relationship exists, that is, a relationship where one individual is a direct descendant of the other (parent, grandparent, great-grandparent, etc.), the MD is the length of the shortest path between the individuals. Otherwise, the MD is the sum of the distances between each individual and their most recent common ancestor.

## Results

### Validation of Simulated Pedigree Structures

Our software can simulate pedigrees based on user-specified sibship size distributions for each generation. We used our pedigree simulator to simulate 10,000 families across five generations to assess our ability to simulate non-uniform sibship sizes. We used sibship size estimates from IPUMs [**33**] to simulate our pedigrees, specifically for census years spanning from 1850 to 1970 (1850, 1880, 1910, 1940, 1970) **[Table 1]**. We estimated the observed mean and standard deviation of sibship size for the 10,000 simulated pedigrees across each generation. [**Table 1**]. The observed simulated sibship sizes are closely aligned with the sibship size means and standard deviations provided as parameters to the simulation. These findings confirm our ability to simulate pedigrees, accommodating changing sibship sizes across successive generations.

**Table 1:** Validation Results of Our Pedigree Structure Simulator. We present the true distributions of sibship sizes inputted into our pedigree simulator alongside the estimated distributions generated by the simulator.

|  | Mean | SD | Num Sibships Simulated | Observed Mean | Observed SD |
| --- | --- | --- | --- | --- | --- |
| Gen 1850 | 3.31 | 2.13 | 10,000 | 3.32 | 2.03 |
| Gen 1880 | 3.02 | 1.95 | 33,169 | 3.04 | 1.86 |
| Gen 1910 | 2.72 | 1.85 | 100,688 | 2.78 | 1.77 |
| Gen 1940 | 2.35 | 1.70 | 280,343 | 2.41 | 1.61 |
| Gen 1970 | 2.30 | 1.43 | 675,309 | 2.33 | 1.40 |

To demonstrate the impact differing sibship size models may have, we compared a two-child model and an empirical sibship size model. The former assumes each pair of parents has two children together, while the latter draws distributions of sibship sizes from the above-mentioned IPUMs census data. Four-generation pedigrees were simulated under both models, with 1,000 pedigrees simulated under the empirical sibship size model to provide a distribution of pedigrees created with our model. **Fig 4a**, illustrates the relative proportions of 11 relationship types across the two models. The empirical sibship size model exhibits a higher proportion in the number of cousins compared to the two-child model due to the creation of more siblings, particularly in earlier generations. **Fig 4b** compares the distribution of the number of cousins between the two models, showing many more cousins under the empirical sibship size model. The number of cousin relationships per individual varies substantially, particularly among more distant cousins. Overall, our pedigree simulator demonstrates the capacity to generate pedigrees with a more realistic representation of distant relationships by offering user-provided varying sibship sizes over generations.

**Fig 4.**
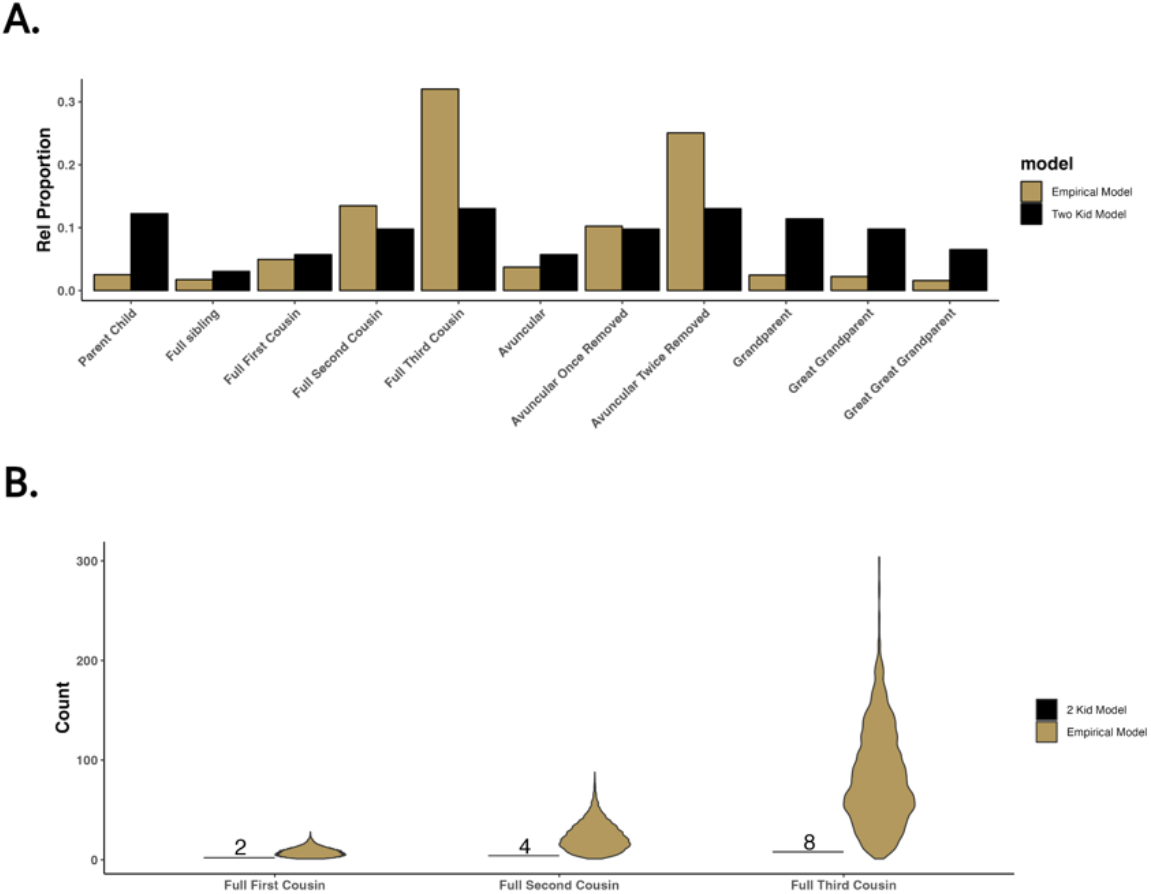
Comparison of two simulated family pedigree models, a two-kid model, and an empirical model. The empirical model is colored in gold, while the 2 kid model is shown in black. A) The relative proportion of eleven relationships found across the two-family models. B) Individual-level variation in the degree of cousin’s relationships across the two models.

### Validation of Genetic Relationships via Estimated Kinship

To test the validity of our software’s SLiM wrapper to simulate genomes onto pedigrees, we compared the estimated kinship coefficient to their expectation for parent-child relationships in six pedigrees with simulated genomes [**Table 2**]. Using our pedigree simulator software, we simulated six families to simulate genomes and estimate their kinship. The six families simulated were made to increase in size for both the number of founders needed to initialize and the total size of the family. We used the kinship estimator Goudet *et al*. proposed due to its ability to provide stable estimates in small samples **[34, 35]**. Founders were initialized using empirical data from the 1000 genomes consortium using individuals from the African superpopulation group.

**Table 2:** Six simulated families used to simulate genomes using our software. To initialize founders with the African Super Population from the 1000 genomes consortium, we had to ensure the families simulated have under 662 founders. Families simulated an increase in size.

|  | Number of Founders | Total Family Size |
| --- | --- | --- |
| Fam 1 | 98 | 726 |
| Fam 2 | 180 | 1287 |
| Fam 3 | 320 | 2277 |
| Fam 4 | 454 | 3392 |
| Fam 5 | 542 | 3555 |
| Fam 6 | 608 | 4021 |

The parent-child relationships for all six families have a kinship estimate within the expectation of 0.25 **[Fig 5]**. While we notice a slight underestimation of kinship, it is noteworthy that the average kinship within each family increased gradually as the family size expanded. However, the observed increase remained relatively modest. One explanation for this downward bias we observe is relatedness within the founders used inside the 1000 genome consortia [**36**]. We estimated kinship for more distant genetic relationships using the largest family simulated (Family 6). The average kinship estimates for all seven relationships converged closely to their expected values in the pedigree. **[Figure 5b] [Table 3]**. Finally, for the sixth family, we looked at the observed vs expected kinship within all genetic relatives inside the pedigree. We estimated the expected kinship with KinInbCoef. We found a strong correlation (R^2 = 0.89) between observed and expected kinship [**Fig 5c**]. These results demonstrate py_ped_sim’s capability to simulate genomes onto family pedigrees with expected levels of kinship across all relationships with expected genetic kinship. This validation also extends to the SLiM simulator, as our software successfully reconstructs pedigrees for SLiM’s simulations.

**Table 3:** Mean observed kinship of the simulated genomes for Family Six across seven different relationship statistics. Each relationship is provided with the expected kinship.

| Relationship | Expected Kinship | Mean Observed Kinship |
| --- | --- | --- |
| Parent-Child | 0.25 | 0.246 |
| Sibling | 0.25 | 0.247 |
| Grandparent | 0.125 | 0.122 |
| Avuncular | 0.125 | 0.123 |
| 1st Cousin | 0.063 | 0.057 |
| Great Grandparent | 0.063 | 0.063 |
| 2nd Cousin | 0.016 | 0.010 |

**Figure 5:**
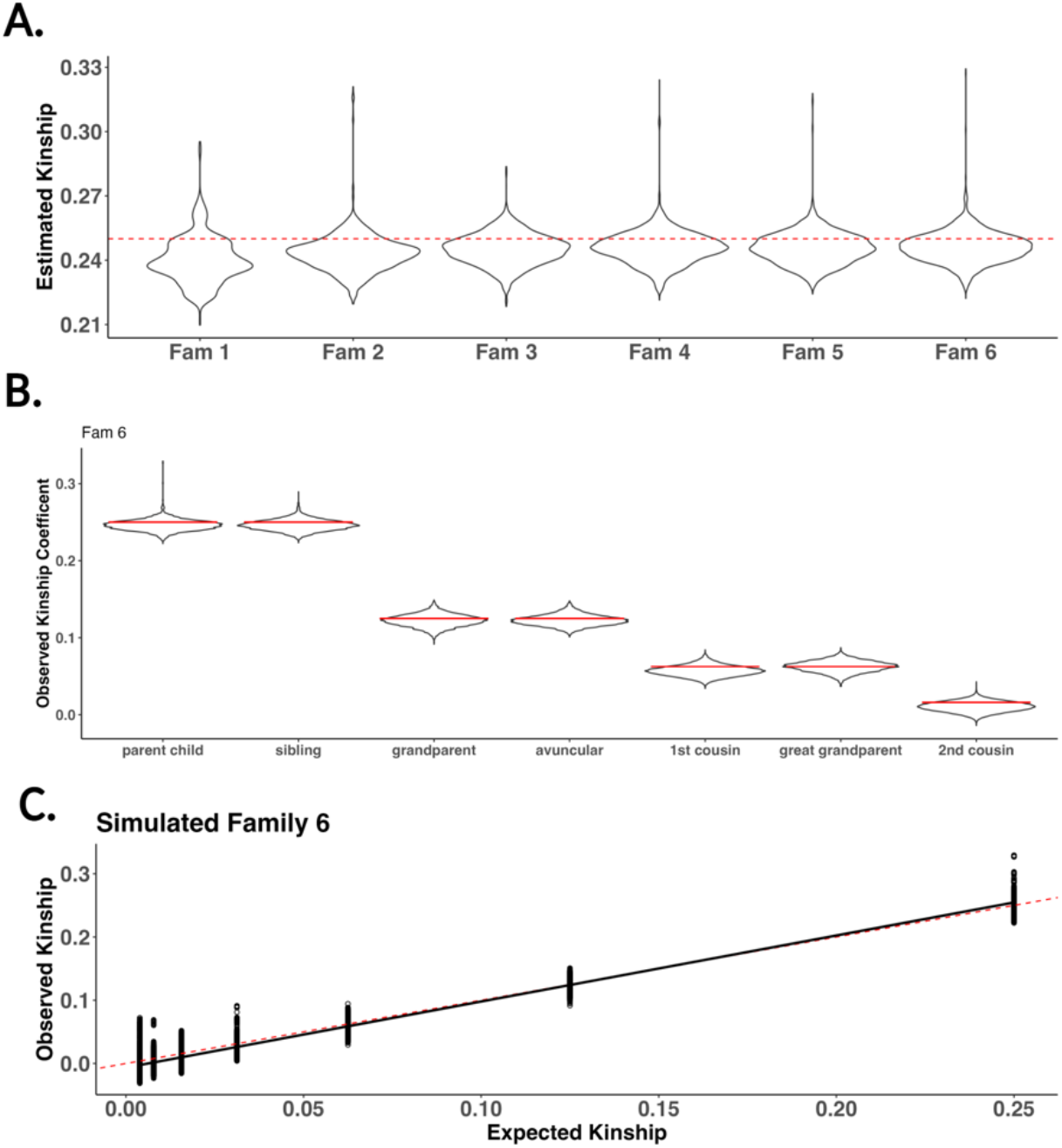
Estimated kinship for six census simulated families using py_ped_sim. (A) We plot the distributions of observed kinship for all parent-child relationships for the six families simulated. (B) For Family 6, the largest family, we observed the estimated kinships for seven relationships found in our family. (C) Scatter plot of observed vs expected kinship for all relationships found in the family pedigree. The solid black line is a regression between estimated and expected kinship. The red dashed line represents the expected kinship across each relationship.

### Kinship Estimations across Various Assumptions of Pedigree Structure

Next, we want to consider how expected vs observed kinship will change across various pedigree structures. We simulate genomes onto five pedigree structures, comparing five different structures of pedigrees. We analyzed five pedigrees: an empirical family obtained from Familinx, a simulated 2-child model, a census-based simulated family, and two simulated families with varying levels of misattributed paternity (MAP=0.01, 0.05) **[Table 4]**. For the simulated 2-kid model, we simulated a pedigree where each set of parents has two kids, not varying the sibship rates across generations. For the five pedigree structures considered, we kept the size of the family (2200-2600 individuals) and number of founders initalized (608-632) the same **[Table 4]**. The only exception to this is the 2-kid simulated family, having a pedigree of 1,542 individuals and a founder size of 512 individuals. This is due to the growth in the deterministic family pedigree model when adding another generation, creating more founders than can be initialized by the African superpopulation.

**Table 4:** Five families used to simulate genomes using our genome simulator software. CS stands for census family pedigrees simulated by our software, with MAP being additional simulations of non-paternity events.

|  | Pedigree Structure Type | Number of Founders | Total Family Size |
| --- | --- | --- | --- |
| Fam 1 | Empirical | 608 | 2198 |
| Fam 2 | 2-Kid Simulated | 512 | 1534 |
| Fam 3 | Census Simulated (CS) | 616 | 2523 |
| Fam 4 | CS + MAP (0.01) | 621 | 2528 |
| Fam 5 | CS + MAP (0.01) | 632 | 2539 |

Expected and observed kinship are strongly correlated for all the pedigree structures **[Figure 6]**. However, the empirical family had incomplete data, including offspring with only one known parent, thus including implicit founders. While our simulation framework allows the ability to simulate genomes for missing parents, the resulting VCF output does not contain the genomic profile of the implicit founder.

**Figure 6:**
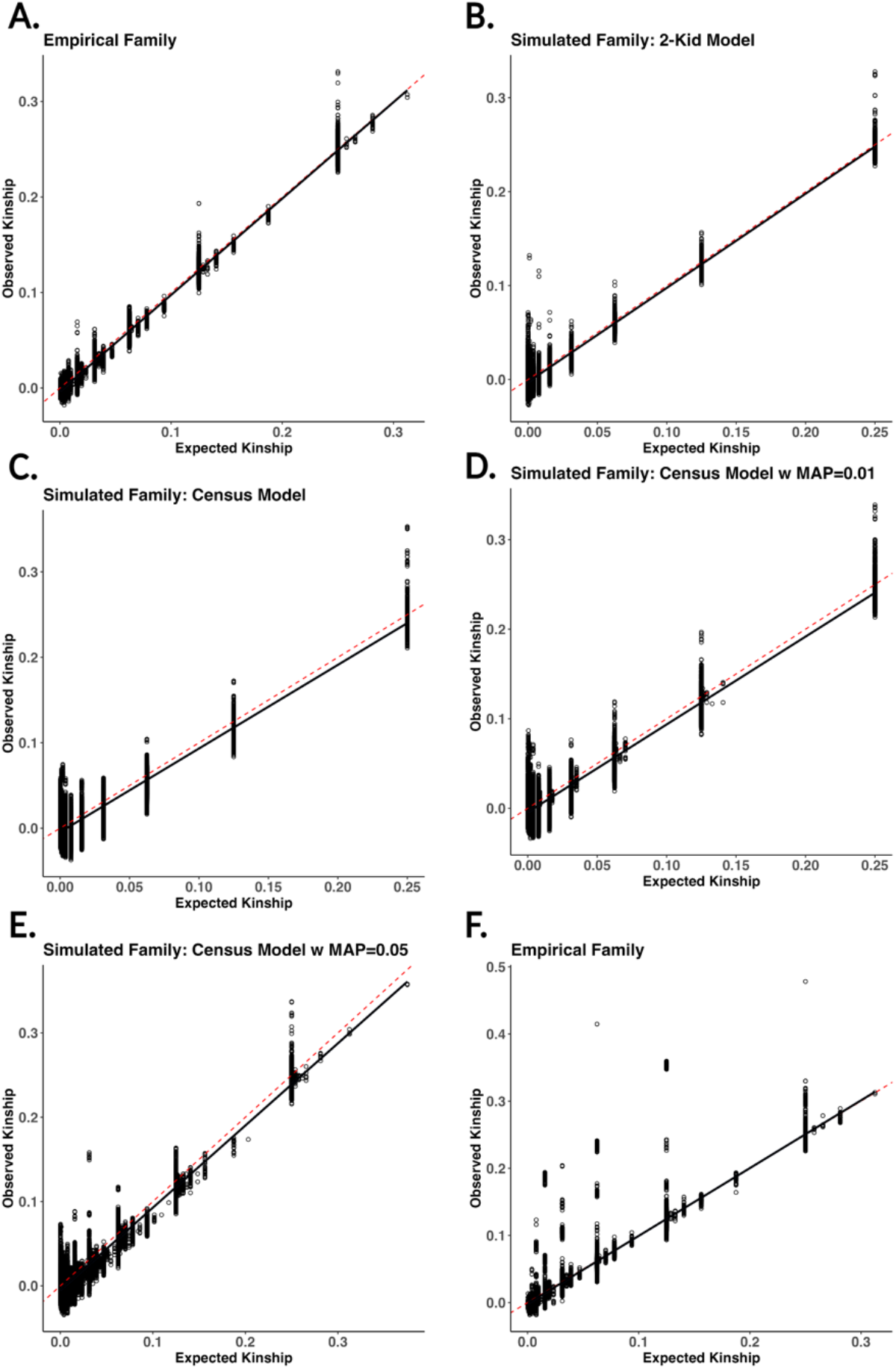
Scatter plots of estimated vs expected kinship across six different pedigree structures. The solid black line is a regression between estimated and expected kinship. The six different pedigrees generated represent A) empirical pedigree from Familinx not including implicit founders, B) simulated family using a 2-kid model, C) simulated family using census data for sibship distributions, D-E) census simulated family with simulated misattributed paternity (MAP = 0.01, 0.05), and F) empirical family from Familinx with implicit founders included. The red dashed line represents the expected kinship across each relationship.

This will introduce bias into the kinship estimate by not accounting for the absent parent’s genetic contribution, which is a property of the kinship estimator itself **[34, 35]**. Repeating the simulation by making all the founders in the pedigree explicit, the best line of fit improved in the empirical family,increasing the r2 value from 0.96 to 0.99 **[Fig 6e]**. Overall, our results underscore the ability of our software to simulate pedigrees with kinship estimates that align with expectations across various pedigree structures.

## Discussion

py_ped_sim offers open-source tools for complex pedigree simulations and facilitates genome simulation onto pedigree structures using SLiM. We present a pedigree simulator that specifies distribution parameters for sibship size across generations, allowing for generational shifts in sibship rates. Additionally, by incorporating genetic half relationships into the simulation of family pedigrees, we further allow the modeling of realistic pedigrees. We offer the ability to simulate genomes based on user-provided pedigree structures. SLiM requires users to include the generation number for each individual and identify which individuals are founders, making it cumbersome for users. Users can create the starting genomes to use for founders, allowing the ability to simulate pedigrees from diverse populations and demographic histories. Finally, we also present a feature to identify genetic relationships between individuals within a family pedigree using three generalizable relationship metrics.

Our validation efforts demonstrate the software’s proficiency in simulating genetic relationships with expected kinship levels and confirming its ability to simulate sibships according to the provided sibship size distributions. This involved assessing the correlation between observed and expected kinship values in simulated pedigrees across various empirical and simulated pedigree structures. We additionally provide validation scenarios for our misattributed paternity and pedigree simulator features. We can simulate the distribution of misattributed paternity events and number of sibships simulated based on user-provided parameters.

Despite these advancements, our software does have limitations. Forward simulations can be computationally demanding, particularly when initializing founders with large genomes. By incorporating tree sequences into genomic simulations in SLiM, run time could be reduced, as they can speed up simulations under feed-forward approaches **[37]**. An important limitation of our pedigree simulator is the ability to simulate different rates of sibship for pairs of parents in the same generation if the user wants to use different sets of sibship distributions across different parts of the pedigree. Another significant limitation is the lack of known half-sib relationships in our simulations. While our approach simulates half-genetic relationships by accounting for misattributed paternity, it does not simulate other scenarios involving half-genetic relationships.

Overall, py_ped_sim will help facilitate genomic analyses based on pedigrees across medical, evolutionary, and forensic sciences. While our primary focus on pedigree simulations centers on human kinship, this approach is adaptable to non-human organisms. This tool will allow simulations of large-scale ecological and evolutionary pedigrees, offering insight into inheritance patterns and understanding the evolutionary dynamics of populations. In forensics, py_ped_sim can be used to explore the performance of Investigative Genetic Genology over pedigree structures and genomic variation. We additionally offer a feature to generalize relationships into multiple metric relationships. Generalizing relationships between pairs of individuals within a family pedigree facilitates identifying connections across extensive family networks and streamlines the process of recognizing relationships across multiple pedigrees.

In conclusion, py_ped_sim provides an accessible solution for accurate pedigree and genome simulations. Forward simulations create genetic relationships across diverse family pedigree structures with varying sibship sizes across generations. This software establishes a basis for future research and paves the way for advancements in genomic studies involving pedigrees.

## Ethics approval and consent to participate

Not Applicable

## Consent for publication

Not Applicable

## Availability and Requirements

Project name: py_ped_sim - A flexible forward genetic simulator for complex family pedigree analysis Project home page: https://github.com/MiguelGuardado/ped_slim Operating System: Windows, MacOS, Linux Programming Language: Python, SLiM for genetic simulations Other requirements: mini-conda for virtual environment dependencies. License: GPL-3.0

Any restrictions to use by non-academics: None

## Competing interests

The authors declare no competing interests.

## Funding

This project, particularly R.V.R., M.G., was supported by NIJ grant 2019-DU-BX-0028. M.G. is a recipient of a Howard Hughes Medical Institute Gilliam Fellowship, Achievement Award for College Scientists Foundation Scholarship, and a UCSF Discovery Fellows Program Award. S.J. and B.C. was supported by the Genentech Foundation Scholars #G-7874540. The NIH MBRS-RISE R25-GM059298 supported J.M and C.P.. E.S. and M.G. were supported by the NIH MARC T34-GM008574. The NIH U-RISE T34-GM145400 supported S.C.. Finally, C.P. was supported by NSF LSAMP HRD-1826490

## Availability of Data and Materials

Software can be found on Git Hub (https://github.com/MiguelGuardado/ped_slim). Empirical founder genomes used from 1000 genomes consortium can be found on their FTP site (https://www.internationalgenome.org/data/).

## Authors’ contributions

Conceptualization – M.G., C.P., R.R.

Software programming – M.G., C.P., S.C., J.M., S.J., B.C. K.S., E.S., S.H.

Formal analysis – M.G., C.P, J.M, S.C., S.J., B.C.

Investigation - M.G., C.P, J.M, S.C., S.J., B.C.

Writing-Manuscript – M.G., E.Z., R.R

Software Testing – M.G., S.C., S.J., J.M., K.S., E.S. Validation – M.G., S.C., E.Z., R.R.

Visualization – S.C., M.G Supervision – R.R., E.Z., R. H. Project administration – R.R., E.Z.

## Acknowledgments

We extend our gratitude to Camila Tamburrini for the valuable discussions and testing conducted to enhance the usability of our software. M.G. is a recipient of a Howard Hughes Medical Institute Gilliam Fellowship, Achievement Award for College Scientists Foundation Scholarship, UCSF Discovery Fellows Program Award, and UCSF Initiative for Maximizing Student Development. E.I.Z. received funding from the Miller Institute for Basic Research in Science, University of California Berkeley. Finally, we thank the team that wrote ‘Equity in Author Order: A Feminist Laboratory’s Approach’ for providing a helpful model for establishing authorship order.

## Notes

### Competing Interest Statement

The authors have declared no competing interest.

## References

1. Li Y-J, Scott WK, Hedges DJ, Zhang F, Gaskell PC, Nance MA, Watts RL, Hubble JP, Koller WC, Pahwa R, Stern MB, Hiner BC, Jankovic J, Allen FA Jr, Goetz CG, Mastaglia F, Stajich JM, Gibson RA, Middleton LT, Saunders AM, Scott BL, Small GW, Nicodemus KK, Reed AD, Schmechel DE, Welsh-Bohmer KA, Conneally PM, Roses AD, Gilbert JR, Vance JM, Haines JL, Pericak-Vance MA: Age at onset in two common neurodegenerative diseases is genetically controlled. Am. J. Hum. Genet. 2002, 70:985–993.

2. Glahn DC, Nimgaonkar VL, Raventós H, Contreras J, McIntosh AM, Thomson PA, Jablensky A, McCarthy NS, Charlesworth JC, Blackburn NB, Peralta JM, Knowles EEM, Mathias SR, Ament SA, McMahon FJ, Gur RC, Bucan M, Curran JE, Almasy L, Gur RE, Blangero J: Rediscovering the value of families for psychiatric genetics research. Mol. Psychiatry 2019, 24:523–535.

3. Kember RL, Hou L, Ji X, Andersen LH, Ghorai A, Estrella LN, Almasy L, McMahon FJ, Brown C, Bućan M: Genetic pleiotropy between mood disorders, metabolic, and endocrine traits in a multigenerational pedigree. Transl. Psychiatry 2018, 8:218.

4. Gibson G: Rare and common variants: twenty arguments. Nat. Rev. Genet. 2012, 13:135–145.

5. McClellan JM, Susser E, King M-C: Schizophrenia: a common disease caused by multiple rare alleles. Br. J. Psychiatry 2007, 190:194–199.

6. Bodmer W, Bonilla C: Common and rare variants in multifactorial susceptibility to common diseases. Nat. Genet. 2008, 40:695–701.

7. Taliun D, Harris DN, Kessler MD, et al.: Sequencing of 53,831 diverse genomes from the NHLBI TOPMed Program. Nature 2021, 590:290–299.

8. Nguyen TN, Chen N, Cosgrove EJ, Bowman R, Fitzpatrick JW, Clark AG: Dynamics of reduced genetic diversity in increasingly fragmented populations of Florida scrub jays,Aphelocoma coerulescens. Evol. Appl. 2022, 15:1018–1027.

9. Fioretti M, Negrini R, Biffani S, Quaglia A, Valentini A, Nardone A: Demographic structure and population dynamics of Maremmana cattle local breed after 35 years of traditional selection. Livest. Sci. 2020, 232:103903.

10. Wilson AJ, Nussey DH, Pemberton JM, Pilkington JG, Morris A, Pelletier F, Clutton-Brock TH, Kruuk LEB: Evidence for a genetic basis of aging in two wild vertebrate populations. Curr. Biol. 2007, 17:2136–2142.

11. Doris PA: The genetics of blood pressure and hypertension: the role of rare variation. Cardiovasc. Ther. 2011, 29:37–45.

12. Umer S, Zhao SJ, Sammad A, Weldegebriall Sahlu B, Yunwei P, Zhu H: AMH: Could It Be Used as A Biomarker for Fertility and Superovulation in Domestic Animals? Genes 2019, 10.

13. Nawaz MY, Jimenez-Krassel F, Steibel JP, Lu Y, Baktula A, Vukasinovic N, Neuder L, Ireland JLH, Ireland JJ, Tempelman RJ: Genomic heritability and genome-wide association analysis of anti-Müllerian hormone in Holstein dairy heifers. J. Dairy Sci. 2018, 101:8063–8075.

14. Grueber CE, Wallis GP, Jamieson IG: Genetic drift outweighs natural selection at toll-like receptor (TLR) immunity loci in a re-introduced population of a threatened species. Mol. Ecol. 2013, 22:4470–4482.

15. Fradgley N, Gardner KA, Cockram J, Elderfield J, Hickey JM, Howell P, Jackson R, Mackay IJ: A large-scale pedigree resource of wheat reveals evidence for adaptation and selection by breeders. PLoS Biol. 2019, 17:e3000071.

16. Kling D, Phillips C, Kennett D, Tillmar A: Investigative genetic genealogy: Current methods, knowledge and practice. Forensic Sci. Int. Genet. 2021, 52:102474.

17. Edge M (doc), Coop G: How lucky was the genetic investigation in the Golden State Killer case? bioRxiv 2019:531384.

18. Kling D, Phillips C, Kennett D, Tillmar A: Investigative genetic genealogy: Current methods, knowledge and practice. Forensic Sci. Int. Genet. 2021, 52:102474.

19. Ge J, Budowle B: Forensic investigation approaches of searching relatives in DNA databases. J. Forensic Sci. 2021, 66:430–443.

20. Katsanis SH: Pedigrees and Perpetrators: Uses of DNA and Genealogy in Forensic Investigations. Annu. Rev. Genomics Hum. Genet. 2020, 21:535–564.

21. Dimitromanolakis A, Xu J, Krol A, Briollais L: sim1000G: a user-friendly genetic variant simulator in R for unrelated individuals and family-based designs. BMC Bioinformatics 2019, 20:26.

22. Li B, Wang GT, Leal SM: Generation of sequence-based data for pedigree-segregating Mendelian or Complex traits. Bioinformatics 2015, 31:3706–3708.

23. Nieuwoudt C, Brooks-Wilson A, Graham J: SimRVSequences: an R package to simulate genetic sequence data for pedigrees. Bioinformatics 2020, 36:2295–2297.

24. Baumdicker F, Bisschop G, Goldstein D, Gower G, Ragsdale AP, Tsambos G, Zhu S, Eldon B, Ellerman EC, Galloway JG, Gladstein AL, Gorjanc G, Guo B, Jeffery B, Kretzschumar WW, Lohse K, Matschiner M, Nelson D, Pope NS, Quinto-Cortés CD, Rodrigues MF, Saunack K, Sellinger T, Thornton K, van Kemenade H, Wohns AW, Wong Y, Gravel S, Kern AD, Koskela J, Ralph PL, Kelleher J: Efficient ancestry and mutation simulation with msprime 1.0. Genetics 2022, 220.

25. Anderson-Trocmé L, Nelson D, Zabad S, Diaz-Papkovich A, Kryukov I, Baya N, Touvier M, Jeffery B, Dina C, Vézina H, Kelleher J, Gravel S: On the genes, genealogies, and geographies of Quebec. Science 2023, 380:849–855.

26. Haller BC, Messer PW: SLiM 4: Multispecies Eco-Evolutionary Modeling. Am. Nat. 2023, 201:E127–E139.

27. Haller BC, Messer PW: SLiM 3: Forward Genetic Simulations Beyond the Wright–Fisher Model. Mol. Biol. Evol. 2019, 36:632–637.

28. Charmantier A, Réale D: How do misassigned paternities affect the estimation of heritability in the wild? Mol. Ecol. 2005, 14:2839–2850.

29. Macintyre S, Sooman A: Non-paternity and prenatal genetic screening. Lancet 1991, 338:869–871.

30. Bellis MA, Hughes K, Hughes S, Ashton JR: Measuring paternal discrepancy and its public health consequences. J. Epidemiol. Community Health 2005, 59:749–754.

31. Präg P: Subjective socio-economic status predicts self-rated health irrespective of objective family socio-economic background. Scand. J. Public Health 2020, 48:707–714.

32. Hagberg A, Swart P S Chult D: Exploring network structure, dynamics, and function using NetworkX. Los Alamos National Lab.(LANL), Los Alamos, NM (United States); 2008.

33. Sarah Flood, Miriam King, Renae Rodgers, Steven Ruggles, J. Robert Warren, Daniel Backman, Annie Chen, Grace Cooper, Stephanie Richards, Megan Schouweiler and Michael Westberry. IPUMS CPS: Version 11.0 [dataset]. Minneapolis, MN: IPUMS, 2023. 10.18128/D030.V11.0.

34. Weir BS, Goudet J: A Unified Characterization of Population Structure and Relatedness. Genetics 2017, 206:2085–2103.

35. Goudet J, Kay T, Weir BS: How to estimate kinship. Mol. Ecol. 2018, 27:4121–4135.

36. 1000 Genomes Project Consortium, Auton A, Brooks LD, Durbin RM, Garrison EP, Kang HM, Korbel JO, Marchini JL, McCarthy S, McVean GA, Abecasis GR: A global reference for human genetic variation. Nature 2015, 526:68–74.

37. Haller BC, Galloway J, Kelleher J, Messer PW, Ralph PL: Tree-sequence recording in SLiM opens new horizons for forward-time simulation of whole genomes. Mol. Ecol. Resour. 2019, 19:552–566.

